# Imaging Double Fertilization in Maize

**DOI:** 10.64898/2026.06.12.731921

**Authors:** Andrea RM. Calhau, Thomas Widiez

## Abstract

Sexual reproduction in flowering plants relies on double fertilization, a process marked by two fusion events between the male and female gametes that lead to seed formation. Because this process unfolds within the embryo sac embedded deep inside the ovule, direct observation remains technically demanding, especially in maize, where the large size of female reproductive organs presents additional obstacles. The described method enables high-resolution visualization of cellular events unfolding during maize double fertilization. The approach integrates optimized fixation, clearing and confocal imaging of embryo sacs from ears pollinated with fluorescent pollen marker lines. Precise timing of embryo sac fixation is critical, allowing capture of key events such as pollen peri-germ cell membrane break-down or gamete karyogamy. The protocol provides detailed guidance for ovule dissection, fixation, preparation and renewal of the clearing solution and confocal imaging of embryo sacs. This method offers unprecedented access to the cellular events of double fertilization in maize, establishing a robust framework for studying reproductive processes and supporting future discoveries in plant reproduction.

## Introduction

Double fertilization can be considered as the beginning of seed development, and is characterized by two fusion events between two male and two female gametes. This unique process is crucial for angiosperm reproduction and requires precise timing as well as a tight molecular and cellular coordination (Zhong et al., 2025). In addition to the identification of signaling pathways involved in plant gamete interactions (Sprunck, 2020; Zhong et al., 2025), a major breakthrough has been the ability to image double fertilization in *Arabidopsis thaliana*, particularly through advances in live-cell imaging (Denninger et al., 2014; Hamamura et al., 2011, 2014; Mao et al., 2023; Mizuta et al., 2024; Wang et al., 2024). Thanks to fluorescent markers expressed in either the male and/or the female gametophyte, imaging methods have been instrumental in visualizing and dissecting the dynamic cellular events during double fertilization. For example, to characterize sperm cell behavior (Hamamura et al., 2011), pollen tube guidance (Mizuta et al., 2024) and the behavior of peri-germ cell membrane (Sugi et al., 2023, 2024). Imaging has also been central to characterization of key molecular players involved in the fertilization processes in *Arabidopsis* (Maruyama et al., 2013; Mori et al., 2014; Wang et al., 2022; Yu et al., 2021).

While double fertilization imaging techniques have been now widely applied in the model organism *Arabidopsis*, their adaptation to other species, particularly crops like maize, remains limited. A major obstacle is the deep embedding of female gametes within maternal tissues, which complicates imaging. In maize, ovules are significantly large (several millimeters, compared to a few micrometers in *Arabidopsis*), further increasing the technical difficulty for visualizing the embryo sacs. To address these challenges, a specialized protocol was developed. The workflow includes optimized procedures for dissecting and fixing maize ovules, preparing and refreshing the clearing solution, and conducting confocal imaging of embryo sacs from ears pollinated with fluorescent pollen marker lines. Notably, the precise choice of the time after pollination is a key parameter to observe the desired cellular events of maize double fertilization. This method thus provides a solid pipeline for investigating maize double fertilization processes.

## Materials

It is essential that you consult the appropriate Material Safety Data Sheets and your institution’s Environmental Health and Safety Office for proper handling of equipment and hazardous materials used in this protocol.

Recipes: Please see the end of this protocol for a list of selected recipes.

### Biological Materials and Reagents

- Freshly prepared D-PBS (Dulbecco’s Phosphate Buffered Saline) 1X (dilution in ultra-pure water of a 10X D-PBS (Dominique Dutscher: Cat number: 509590)
- Freshly prepared 4% Paraformaldehyde solution (dilution in 1X D-PBS of a 16% PFA solution, Electronic Microscopy Sciences, cat number: 15710)
- ClearSeeAlpha (see “Recipes” section)
- 0.1% Calclofluor White (Fluorescent Brightener 28) solution prepared in ClearSee (Sigma Aldrich, cat number: F3543, CAS number: 4404-43-7)

### Equipment and supplies

- Scissors
- Curved tweezers
- Square petri dish
- Beaker
- Razor blades (Seedburo Equipment Company, seller reference: 121-3)
- Scalpel
- Binocular loupe, called stereomicroscope hereafter.
- Fume hood
- Basic shaker: IKA rocker 2D digital (Dominique Dutscher, seller reference: 250277)
- MAC Smart Strainer 100µm (Miltenyi Biotec, seller reference 130-098-463)
- 50ml Falcon tubes
- 50ml Flacon holder
- Weighing scale
- 26 x 76 mm microscopy slides (Dominique Dutscher, seller reference: 068763)
- 22 x 60 mm coverslip (Dominique Dutscher, seller reference 100267)
- DuPont MOLYKOTE® High Vacuum Grease, 5.3 oz (Cole-Parmer, seller reference 7975130)
- 5mL syringe
- Pen Liquid Blocker: Liquid Blocker Super PAP pen (Fisher scientific, catalogue No. NC9827128)
- ZEISS AxioImager M2 microscope with a dry 10x objective (EC Plan –NEOFLUAR 10x/0.3a ZEISS).
- SP8 laser scanning confocal upright microscope (DM6000) with a water immersion objective (HC FLUOTAR L 25x/0.95 W, from Leica, Wetzlar, Germany) (LEICA)

## Methods

The objective of this protocol is the visualization of cellular events occurring during maize double fertilization. By using pollen from transgenic lines expressing one or multiple fluorescent marker of interest, it allows to monitor distinct cellular compartments throughout the fertilization process. Some potential transgenic lines of interest include lines expressing the chromatin marker Histone 2B (H2B), marking nuclear DNA in the vegetative nucleus and/or sperm cells nuclei (Gilles et al., 2021; Howe et al., 2012), as well as a α-tubulin-YFP line, which labels microtubules in the generative cell and sperm cells (Kliwer & Dresselhaus, 2010; Li et al., 2024). In addition, a NOT-LIKE-DAD translational fusion with the fluorescent protein mCitrine (NLD:mCitrine) can also be used in this protocol as it labels the peri-germ cell membrane (PGCM), an internal membrane of the pollen vegetative cell that surrounds the generative cell sperm cells at early developmental stages and later enclose the two sperm cells (Gilles et al., 2021; Sugi et al., 2024). Confocal imaging of the maize ovule is made feasible through a combination of dissection, fixation, and clearing steps.

### Pollination of a maize ear with pollen from a fluorescent marker line

1. Select an ear suitable for pollination. For the A188 inbred line, the ear should have abundant silks measuring approximately 8-15cm in length (**Fig. 1A**).

> *Depending on the cellular events to be analysed, use pollen from an appropriate transgenic maize line expressing the fluorescent marker of interest. In the following examples, the protocol was applied using two different fluorescent pollen for two purposes: 1) to track the fate of the pollen peri-germ cell membrane after pollen tube discharge in the embryo sac, which was feasible thanks to a peri-germ cell membrane marker line (NLD:mCitrine); 2) to visualize interactions between the nuclei of the male and female gametes, using pollen with fluorescent sperm cell nuclei (See Discussion for further information)*.

2. Schedule the pollination according to the objectives of the experiment and the specific growth conditions.

> *Our growing condition are 16-h illumination period at 25/19°C [day/night]. To assess the fate of the peri-germ cell membrane, pollinate the ears about 18-23h hour before dissection, whereas to evaluate gamete karyogamy nuclei perform the pollination about 26-29h before dissection (See Discussion for further information)*.

**Figure 1:**
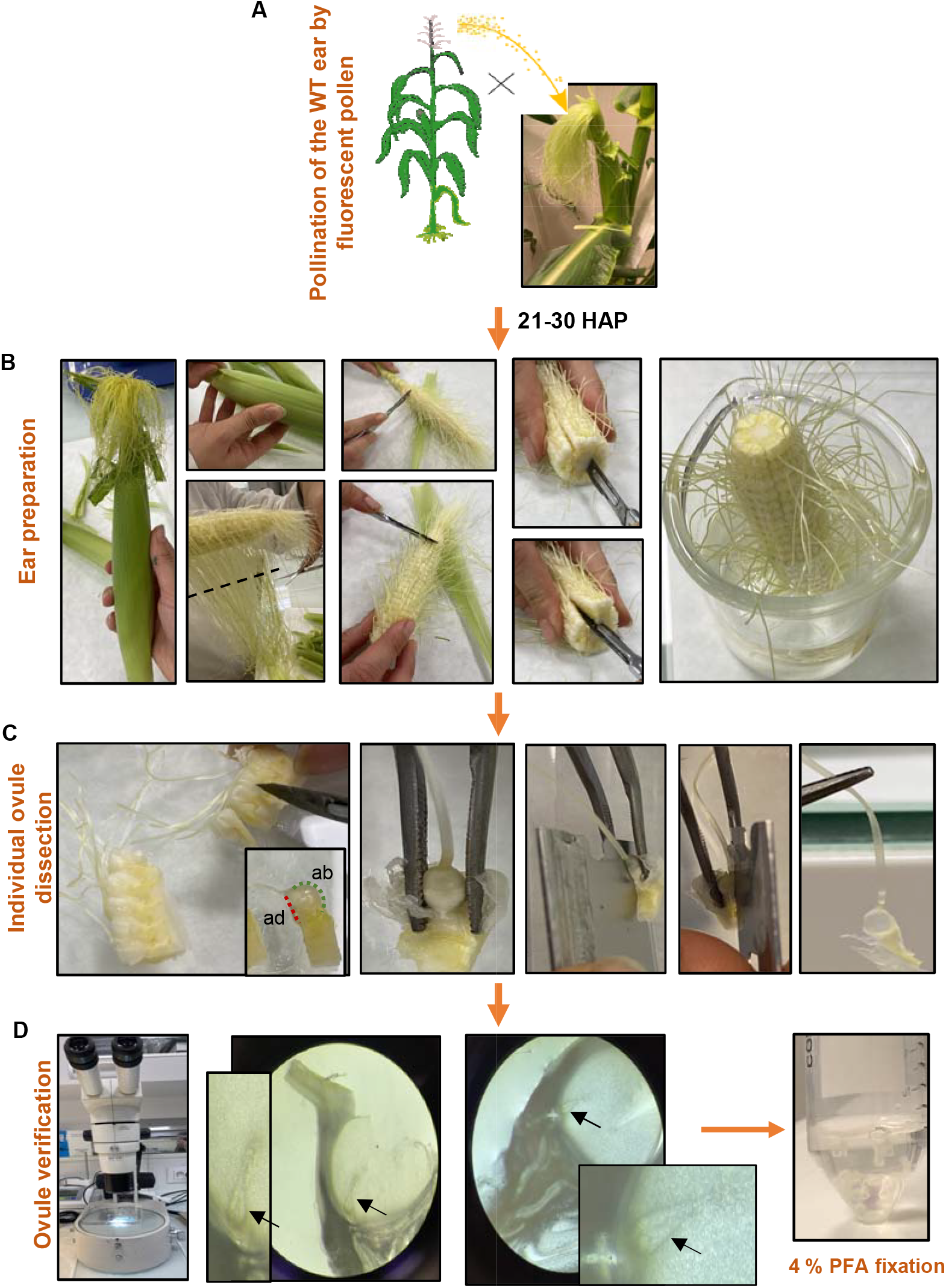
Detailed gestures for ovule dissection. **A)** Illustration of the pollination of WT ear with a fluorescent pollen marker line. **B)** Preparation for ovule isolation. The dashed line indicates the trajectory of the scissors used to cut the silks **C)** Illustration of individual ovule dissection. In the first panel, the red dotted line shows the ventral side (adaxial, ad) and the green dotted line shows the rounded dorsal (abaxial, ab) side. **D)** Observation of embryo sacs (arrows) under a stereo-microscope after dissection and subsequent PFA fixation.

### Dissection of the ovules and Paraformaldehyde (PFA) fixation

3. On the day of the dissection, start by preparing your 4% PFA (Paraformaldehyde) solution.

> *Because pollen tube growth continues after ovule harvest, divide your dissections into separate tubes to distinguish early from late harvested ovules by making hourly timepoints. To do this, split the 4% PFA solution into several 50 mL Falcon tubes. For example, for a 4-hour dissection, 4 hourly timepoints are done so prepare four tubes each containing 10 mL of 4% PFA. Place your harvested ovules in a tube and start filling a new tube with your dissected ovules every hour. Keep the tubes containing the 4% PFA constantly on ice an under the fume hood*.

4. Harvest the ear at the chosen time after pollination by cutting its base and take it to the location of the dissection. Slowly and carefully, remove the husk leaves from the ear (**Fig. 1B**).

> *Try not to pull on or to ruin the silks while removing the husk leaves, since it could damage the ovules*.

5. Cut the silks with the scissors, leaving 1 to 5cm to facilitate later ovule dissection (**Fig. 1B**).
6. To isolate rows of 5 to 10 ovules that remain attached to the maternal tissues of the ear, use a scalpel blade to make an incision on each side of the ovules (**Fig. 1B**), followed by an incision underneath them (**Fig. 1B**). After finishing cutting and removing your row of ovules, put the rest of the ear in a beaker that contains tap water to avoid dehydration of the other ovules.
7. Place the row of ovules on an inverted Petri dish, which will serve as a cutting surface. Carefully cut between the ovules to separate them individually (**Fig. 1C**), taking care not to damage the 1– 5 cm silks attached to each ovule.
8. Cutting both sides of the ovule is necessary to allow efficient fixative penetration (see step 7) and above all, to access the embryo sac for imaging. Therefore, each ovule must be dissected individually and carefully. Ovules have a flat ventral side (adaxial, red dotted line **Fig. 1C**) and a rounded dorsal (abaxial, green dotted line **Fig. 1C**) side. Carefully remove the maternal tissues (glumes) covering the ovule (this can be done with a fingernail) while holding the ovule steady with a pair of curved tweezers. Place the ovule with its flat ventral side facing upward (**Fig. 1C**). Using one hand to stabilize the ovule/tweezer setup, use the other hand to make cuts on both lateral sides of the silk with a razor blade, using the silk as a guide (**Fig. 1C**). After each ovule dissection, check under a stereo-microscope with transmitted light to ensure that the embryo sac is present in the dissected ovule and to assure that section is intact and not damaged (**Fig. 1D**).

> *During this process, the razor blade should be slightly tilted so that the abaxial side becomes thinner when considering the final shape of the section containing the female gametophyte (see* ***Fig. 1C****). The razor blade needs to be sharp to avoid crushing the tissues. If the razor becomes blunt, change it*.

9. Cut to shorten the silk, leaving approximately 0.5-1cm of length, as it will help later during the preparation of the slides. Put the dissected ovule on the 4% PFA solution in a 50ml falcon tube that is under the fume hood and on ice.

> *If the fume hood is a bit far away from the bench, you can alternatively collect multiple dissected ovules in 1X D-PBS and transfer them together to the 4% PFA solution. However, avoid prolonged time in 1X D-PBS before fixation to prevent sample degradation*.

10. At the end of the dissection, replenish the ice supporting the tubes containing 4% PFA and the dissected ovules. Place the box of ice with opened tubes under a vacuum bell chamber. Vacuum infiltration, enhances PFA absorbance into the tissues.

> *⚠Tubes must be open. Leave the lids on the side*.

11. Close the vacuum bell chamber when it reaches -0.06 mPa and maintain the vacuum for 15 min. Then, gradually release the vacuum. Repeat this process five times.

> *⚠Release the vacuum extremely slowly to not damage the cells*.

12. Renew the 4% PFA solution with a fresh 4% PFA solution, wrap the tubes in aluminium foil and incubate overnight at 4°C.

> *MAC Smart Strainer 100µm can be used to change the 4% PFA solution as a sort of “sieve” to retain the samples*.
>
> *This is toxic waste. Dispose of it accordingly*.
>
> *The aluminium foil keeps samples from light that could damage fluorescent proteins*.

13. If fixation is successful, the samples will settle at the bottom of the falcon tube. Rinse samples three times with 1X D-PBS, 1 min each.

> *MAC Smart Strainer 100µm can be used as a sort of “sieve” to retain the samples while changing your rinsing baths*.
>
> *This is toxic waste. Dispose of it accordingly*.

### Clearing of the samples and calcofluor staining

*To visualize the inside of maize embryo sacs, samples must undergo a clearing process. Use ClearSeeAlpha, as it is compatible with fluorescent imaging* (Kurihara et al., 2021).

14. Start by preparing the required volume of ClearSeeAlpha solution (refer to the “Recipes” section).
15. Immediately after the last PBS rinse, transfer the content of each 50mL falcon to a new one.

> *To do this transfer, add ∼20 ml of 1X D-PBS on the original tube, agitate and pour its content into the new tube*.

16. Once all samples are in the new tube, discard the 1X D-PBS and replace it with ClearSeeAlpha to start the clearing of the tissues.

> *MAC Smart Strainer 100µm can be used as a sort of “sieve” to retain the samples while changing from 1X D-PBS to ClearSeeAlpha*.
>
> *The volume of ClearSeeAlpha required depends on the number of samples; ensure all samples are fully covered in the ClearSeeAlpha solution*.

17. Incubate the tubes at room temperature with gentle agitation, e.g., using the basic shaker “IKA rocker 2D digital” and setting speed at 15. Wrap the tubes in aluminium foil.

> *Samples must remain under soft agitation until they are completely cleared*.

18. Renew the ClearSeeAlpha solution 2-3 times per week until samples are cleared.

> *The ClearSeeAlpha solution has to be freshly prepared (see recipes)*.
>
> *The clearing time varies depending of the samples. Since they are manually sectioned the duration depends on their thickness and on the position of the embryo sac. In our case, clearing has been observed to take 2 weeks up to 2 months*.
>
> *This is toxic waste. Dispose of it accordingly*.

19. Once samples are fully cleared, stain them using a 0.1 % Calcofluor White solution. Remove the ClearSeeAlpha from each 50mL tube and replace it with 0.1% Calcofluor White solution prepared in ClearSee (not ClearSeeAlpha). Incubate at room temperature for 30 minutes.

> *Counterstaining with Calcofluor White is optional, as it only labels maternal tissues around the embryo sac, not the embryo sac per se* (**Fig. 2A**). *However, it is recommended as it facilitates embryo sac localization during confocal observations*.

20. Remove the staining solution from all tubes and replace it with fresh ClearSee solution (not ClearSeeAlpha). Incubate under gentle agitation for 30 minutes.
21. Store your stained samples in ClearSee solution at 4°C.

**Figure 2:**
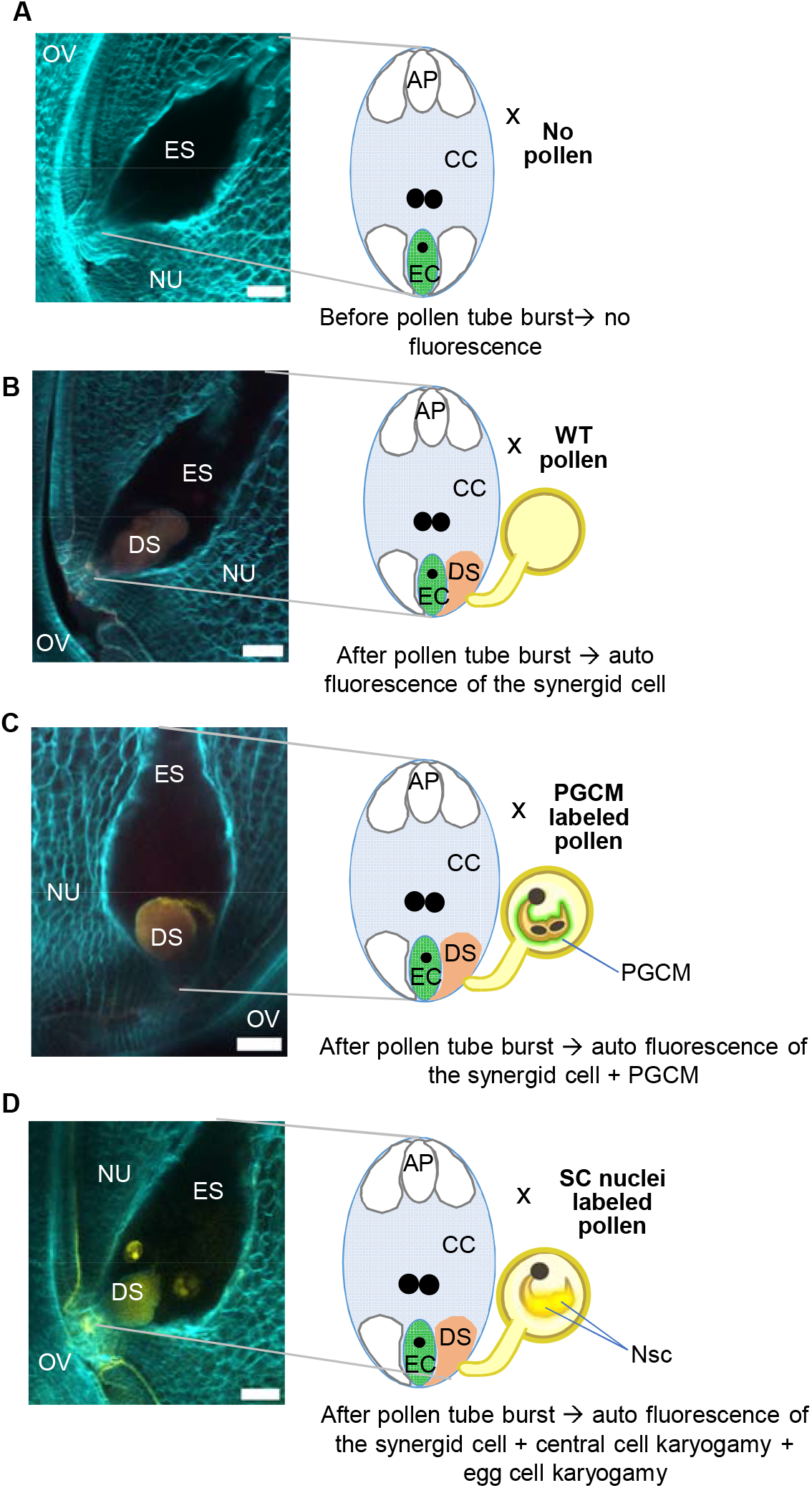
Fluorescent imaging of maize embryo sacs before and after pollen tube burst. **A)** Maize embryo sac prior to pollen tube burst. This situation is observed when no pollen is deposited on the silk (control) or when no pollen tube reaches the observed embryo sac. **B)** Maize embryo sac following wild-type (WT) pollen tube burst. This serves as a negative control to visualize the natural autofluorescence of the degenerating synergid cell. **C)** Maize embryo sac after the burst of a pollen tube containing a fluorescent peri-germ cell membrane (PGCM). The pollen tube burst event is detected by the autofluorescence of the degenerating synergid cell, while the PGCM signal is shown in yellow. **D)** Maize embryo sac after the burst of a pollen tube expressing a sperm cell nuclei marker. The pollen tube burst event is detected by the autofluorescence of the degenerating synergid cell, double fertilization is monitored thanks to the fluorescent chromatin from the male. The presence of two fluorescent nuclei indicates egg cell/sperm cell nuclei karyogamy, whereas three fluorescent signals indicate the central cell/sperm cell karyogamy. AP, antipodal cells; CC, central cell; EC, egg cell; DS, degenerating synergid; ES, embryo sac; NU, nucellus; PGCM, peri-germ cell membrane; Nsc, nuclei sperm cells Scale bar =50µm

### Samples preparation and Confocal observations

#### Microscopic slide preparation

*Once samples are cleared and stained, mount the slides for observation. As the ovules are manually sectioned, thickness variability occurs. Vacuum grease serves as a suspension medium for mounting, facilitating adaptation of the system to varying sample dimensions*.

22. In a microscopy slide, draw a rectangle with a Liquid Blocker pen.

> *Dimensions of the rectangle depend on the number of ovules intended for mounting. In our hands, a rectangle of roughly 3*.*5 x 1*.*5 cm can accommodate 8 to 11 ovules*.
>
> *The liquid blocker pen creates a hydrophobic barrier on the base of the microscopy slide and provides a reference for drawing the rectangle with the high vacuum grease (see below)*.

23. Fill a 5ml syringe with vacuum grease by removing the syringe plunger and pouring the grease directly from the tube into the barrel.
24. Trim the tip of a pipet to enlarge the opening. Attach the modified tip to the syringe that contains the vacuum grease. While securing both components manually, dispense the grease to outline a rectangle above the area demarcated by Liquid Blocker pen.
25. After completing the grease rectangle, gently press the cut tip against the grease to ensure adhesion to the microscopic slide and to prevent ClearSee leakage.

#### Samples mounting

Given the manual sectioning of ovules, the embryo sac position varies within each ovule. To optimize embryo sac visualization, preliminary examination under a visible light microscope is essential for identifying the side offering the clearest view of the embryo sac.

26. Screen the samples using a visible light microscope to evaluate the following criteria: (1) Presence of the embryo sac; (2) Visibility of the embryo sac from both sides of the dissected ovule, selecting the side with optimal clarity for further imaging; (3) On the selected side, assess whether the embryo sac nuclei are distinguishable, and check for potential obscuration of the embryo sac base by maternal tissues. Ovules exhibiting such obscuration should be excluded to prevent compromised confocal imaging.

> *Perform this procedure in a darkened room to minimize light exposure. The use of a DIC filter of a ZEISS AxioImager M2 microscope with a dry 10x objective (EC Plan –NEOFLUAR 10x/0*.*3a ZEISS) is well-suited to this step*.

27. Position the ovules on the prepared slide with the side offering optimal embryo sac visibility facing upward. Repeat this process until the grease rectangle is fully occupied with ovules.

> *The number of dissected ovules accommodated depends on the dimension of the grease rectangle. In our setup, 8-11 ovules can be arranged*.
>
> *Align all ovules in a single row to facilitate acquisition*.
>
> *To minimize light exposure, store the grease-mounted slide in an opaque box, opening it only as needed*.

28. Once all samples are positioned on the microscopy slide, cover them with ClearSee and apply a coverslip.

#### Confocal observations

Observations are conducted using a confocal microscope. A dedicated channel for calcofluor facilitates identification of the embryo sac, while separate channels are assigned to each fluorescent marker under investigation during double fertilization.

> *A Leica SP8 laser scanning confocal upright microscope (DM6000) with a water immersion objective (HC FLUOTAR L 25×/0*.*95 W, Leica, Wetzlar, Germany) captured the images using three channels: Calcofluor (excitation: 405 nm; detection: 413–504 nm), mCitrine (excitation: 514 nm; detection: 520–600 nm), Autofluorescence (excitation: 552 nm; detection: 564-660 nm). The autofluorescence channel was included to detect signals from the exploded synergid and potential autofluorescence of female nuclei*.

#### Image processing and analysis

The images could be processed with Fiji (https://fiji.sc/) (Schindelin et al., 2012) to adjust brightness and contrast, and to create maximum intensity projections.

## Discussion

### Imaging of Cellular Events within the Maize Embryo Sac

This detailed protocol provides a robust framework for visualizing double fertilization events in maize using fluorescent markers carried by pollen. By enabling high resolution observation of embryo sacs, it allows to capture cellular aspects that were previously difficult to resolve. A previous study employed apropionic acid/ethanol mixture for ovule fixation and subsequently cleared the tissue using methyl salicylate, followed by differential Interference Contrast microscopy (Mòl et al., 1994). Although effective for observation of maize gamete fusion, this approach is incompatible with fluorescence imaging. In contrast, the protocol described here (combining PFA fixation with ClearSeeAlpha clearing) preserves fluorescent proteins and supports high-quality fluorescence microscopy. This enables deeper optical penetration depth, more accurate visualization of internal structures, and substantially improved image resolution. Gilles et al., 2021 applied this imaging strategy to monitor cellular dynamics of the peri-germ cell membrane (PGCM, previously referred as the endo-plasma membrane in Gilles et al., 2021) immediately before double fertilization (Gilles et al., 2021). The PGCM is an internal membrane of the pollen vegetative cell that encloses the sperm cells (Sugi et al., 2024). Using pollen expressing fluorescently labeled PGCM coupled with this method, the authors provided the first documentation, for the first time, of the rapid breakdown of the PGCM upon its arrival in the maize female gametophyte (Gilles et al., 2021) (**Figure 2C**). PGCM breakdown enables the release of the sperm cells and has since been corroborated in *Arabidopsis thaliana* through live-imaging analyses (Sugi et al., 2023). Applying the same methodology, but using pollen grains with fluorescently labeled sperm-cell nuclei, makes it possible to visualize gamete karyogamy in both the central cell and the egg cell (**Figure 2D**).

### Appropriate Controls for Accurate Interpretation of Results

To ensure proper interpretation of the images, at least two essential controls are required. First, imaging ovules from a non-pollinated ear allows the observer to become familiar with an unfertilized embryo sac under the experimental conditions (**Figure 2A**). Second, ovules collected from ears pollinated with wild-type pollen provide a reference embryo sac in which pollen tube rupture has occurred. Indeed, upon pollen tube rupture within the synergid cell, this cell become strongly autofluorescent (**Figure 2B**) (Gilles et al., 2021; Leydon et al., 2015). The autofluorescence of the degenerating synergid cell thus serves as a reliable proxy for successful pollen tube arrival and rupture, while also providing a baseline for background autofluorescence in the absence of fluorescent reporters. In addition, this control could be used to calibrate the time needed for the pollen tube to reach the embryo sac, since this depends of the pollen genotype used, the female genotype and most all the growing condition (especially temperature).

### Technical Constraints, and Perspectives for Maize Embryo Sac Imaging

The large size of the maize ovule, which is an important obstacle for the embryo sacs imaging, was solved in these methods by ovule dissection (**Figure 1C, 1D**). In contrast to *Arabidopsis*, where ovules can be imaged intact without dissection due to their smaller size (Desnoyer & Grossniklaus, 2023; Hamamura et al., 2014; Sugi et al., 2023), maize embryo sac locations vary considerably in depth between samples, since they are generated through manual cutting. This variability necessitates individual adjustments of imaging parameters such as gain, laser intensity, making fluorescence quantification challenging. For this reason, this pipeline is primarily suited for qualitative phenotyping of fertilization events. The two examples shown in **Figure 2C and 2D** illustrate the versatility of the protocol across two applications, PGCM breakdown and gametes karyogamy respectively. However, adapting this protocol to different biological questions requires attention to two critical parameters. First, the selection of the fluorescent marker must be aligned with the specific process under investigation. Second, the timing of tissue dissection must be carefully adjusted to ensure that the relevant cellular events could be captured. In the two examples presented here, dissection times were optimized accordingly: (1) to examine PGCM behavior within the embryo sac (**Figure 2C**), ovules were harvested shortly after pollen tube rupture, typically 18–23 hours after pollination under our conditions; and (2) to visualize gamete karyogamy (**Figure 2D**), samples were collected later, around 26–29 hours after pollination. Lastly, although the examples presented here rely on fluorescence originating from the male gametophyte, this protocol is equally applicable when the fluorescent signal is expressed in the female gametophyte. Furthermore, combining these marker lines with maize mutants defective in fertilization, such as haploid-inducer lines (Gilles et al., 2017; Jacquier et al., 2020), could further expand the range of applications. These strategies would allow a more comprehensive and detailed exploration of the fundamental processes driving maize reproduction.

## Acknowledgments

We acknowledge Sophie Boeuf and Frédérique Rozier for technical advices on ClearSeeAlpha preparation, and ovules dissection respectively, Claire Lionnet for precious guidance on microscopy imaging, Justin Berger and Camile Knaupp for technical assistance with maize culture, Hervé Leyral, Isabelle Desbouchages, for technical assistance in the lab, and Cindy Vial, Julie Prata, Laureen Grangier and Nelly Camilleri for administrative assistance. We also are also grateful to Angèle Noh for helping taking the pictures in figure 1, and Naoya Sugi for his critical reading and constructive suggestions on the manuscript. We acknowledge the contribution of SFR Biosciences (Université Claude Bernard Lyon 1, CNRS UAR3444, Inserm US8, ENS de Lyon) Lymic-PLATIM, member of the national infrastructure France-BioImaging, supported by the French National Research Agency (ANR-10-INBS-04) and the help of the staff for assistance during imaging. This work was supported by the French National Research Agency (ANR) (ANR-19-CE20-0012 to T.W.). TW was also supported by ENS de Lyon and from the Plant Biology and Breeding department of INRAE (BAP). A.R.M.C. is currently supported by a PhD fellowship from the Association Nationale de la Recherche et de la Technologie (ANRT) (Grant Cifre no. 2023/1284).

## Author Contributions

Conceptualization: A.R.M.C., T.W. Investigation: A.R.M.C. Visualization: A.R.M.C Writing—original draft: A.R.M.C. Writing—review and editing: A.R.M.C., T.W. Project Administration: T.W. Funding acquisition: T.W. Supervision: T.W.

## Legends for Figures

### Recipes

- *ClearseeAlpha (to be prepared in advance for storage)*

| Product name | Chemical formula | Chemical Abstracts Service number (Cas #) | Quantity for 50mL in Ultra-Pure water (g) | Quantity for 100mL in Ultra-Pure water (g) | Seller and cat number |
| --- | --- | --- | --- | --- | --- |
| Sodium Deoxycholate (BioXtra, ≥98.0%) | $C_{24}H_{39}NaO_4$ | 302-95-4 | 7,5 | 15 | Sigma Aldrich; cat number 30970 |
| Xylitol | $C_5H_{12}O_5$ | 87-99-0 | 5 | 10 | Sigma Aldrich; cat number W507930 |
| Urea | $CH_4N_2O$ | 57-13-6 | 12,5 | 25 | Euromedex; cat number: EU0014-B |
| *Sodium sulfite | $Na_2SO_3$ | 7757-83-7 | *0,315 | *0,63 | Sigma Aldrich; cat number: 239321 |

\* The main difference between ClearSee (Kurihara et al., 2015) and ClearSeeAlpha (Kurihara et al., 2021) is that ClearSeeAlpha requires the addition of a reducing agent, in this case sodium sulfite. Therefore, before each use, add the appropriate amount of sodium sulfite to prepare the ClearSeeAlpha solution. Note: the quantity of sodium sulfite must be adjusted according to the required volume each time.

The ClearSee solution (before addition of the sodium sulfite), can be stored at room temperature.

**Tips for preparing the ClearSee Solution:**

- The sodium Deoxycholate is an irritant when in form of powder. Consequently, it needs to be weighted either under the fume hood or in a sealed container. For example, to prepare 100mL of ClearSee solution, use a 100mL transparent container with a secure lid. After taring the container and the lid, place them under a fume hood and **slowly** add the required amount of sodium deoxycholate. If your weighing scale is not located under the fume hood, temporarily close the container with its lid, weigh the sodium deoxycholate outside the hood under its closed container to reach the desired mass, then immediately return the closed container to the fume hood. Once reaching the desired weight, add a magnetic stir bar **and a small volume of ultra-pure water** (just enough to initiate dissolution of the powder).
- Put the closed container under agitation to start the solubilisation of the deoxycholate.
- Next, weigh the remaining products and add them to the sodium Deoxycholate solution under the fume hood. After the final component is added, add a small additional volume of water to facilitate dissolution, but do not exceed the final volume required. Continue stirring until all powders are completely dissolved, which may require 30 min to 2h of agitation for 50mL.
- Once the solution becomes clear and homogeneous, measure its volume using a graduated cylinder. Adjust the final concentration by adding water to reach the desired final volume. **⚠ Note: The solution is highly viscous**. Before completing the final volume adjustment, rinse the original preparation container with a small volume of water, mix thoroughly to recover any remaining solution, and then transfer the rinse into the measuring cylinder to ensure accurate volume and concentration.

## References

Denninger, P., Bleckmann, A., Lausser, A., Vogler, F., Ott, T., Ehrhardt, D. W., Frommer, W. B., Sprunck, S., Dresselhaus, T., & Grossmann, G. (2014). Male–female communication triggers calcium signatures during fertilization in Arabidopsis. Nature Communications, 5(1), 4645. 10.1038/ncomms5645

Desnoyer, N. J., & Grossniklaus, U. (2023). Live Imaging of Arabidopsis Pollen Tube Reception and Double Fertilization Using the Semi-In Vitro Cum Septum Method. Journal of Visualized Experiments: JoVE, 192. 10.3791/65156

Gilles, L. M., Calhau, A. R. M., La Padula, V., Jacquier, N. M. A., Lionnet, C., Martinant, J.-P., Rogowsky, P. M., & Widiez, T. (2021). Lipid anchoring and electrostatic interactions target NOT-LIKE-DAD to pollen endo-plasma membrane. Journal of Cell Biology, 220(10), e202010077. 10.1083/jcb.202010077

Gilles, L. M., Khaled, A., Laffaire, J., Chaignon, S., Gendrot, G., Laplaige, J., Bergès, H., Beydon, G., Bayle, V., Barret, P., Comadran, J., Martinant, J., Rogowsky, P. M., & Widiez, T. (2017). Loss of pollen-specific phospholipase NOT LIKE DAD triggers gynogenesis in maize. The EMBO Journal, 36(6), 707–717. 10.15252/embj.201796603

Hamamura, Y., Nishimaki, M., Takeuchi, H., Geitmann, A., Kurihara, D., & Higashiyama, T. (2014). Live imaging of calcium spikes during double fertilization in Arabidopsis. Nature Communications, 5(1), 4722. 10.1038/ncomms5722

Hamamura, Y., Saito, C., Awai, C., Kurihara, D., Miyawaki, A., Nakagawa, T., Kanaoka, M. M., Sasaki, N., Nakano, A., Berger, F., & Higashiyama, T. (2011). Live-Cell Imaging Reveals the Dynamics of Two Sperm Cells during Double Fertilization in Arabidopsis thaliana. Current Biology, 21(6), 497–502. 10.1016/j.cub.2011.02.013

Howe, E. S., Clemente, T. E., & Bass, H. W. (2012). Maize Histone H2B-mCherry : A New Fluorescent Chromatin Marker for Somatic and Meiotic Chromosome Research. DNA and Cell Biology, 31(6), 925–938. 10.1089/dna.2011.1514

Jacquier, N. M. A., Gilles, L. M., Pyott, D. E., Martinant, J.-P., Rogowsky, P. M., & Widiez, T. (2020). Puzzling out plant reproduction by haploid induction for innovations in plant breeding. Nature Plants, 6(6), 610–619. 10.1038/s41477-020-0664-9

Kliwer, I., & Dresselhaus, T. (2010). Establishment of the male germline and sperm cell movement during pollen germination and tube growth in maize. Plant Signaling & Behavior, 5(7), 885–889.

Kurihara, D., Mizuta, Y., Nagahara, S., & Higashiyama, T. (2021). ClearSeeAlpha : Advanced Optical Clearing for Whole-Plant Imaging. Plant and Cell Physiology, 62(8), 1302–1310. 10.1093/pcp/pcab033

Kurihara, D., Mizuta, Y., Sato, Y., & Higashiyama, T. (2015). ClearSee : A rapid optical clearing reagent for whole-plant fluorescence imaging. Development, 142(23), 4168–4179. 10.1242/dev.127613

Leydon, A. R., Tsukamoto, T., Dunatunga, D., Qin, Y., Johnson, M. A., & Palanivelu, R. (2015). Pollen Tube Discharge Completes the Process of Synergid Degeneration That Is Initiated by Pollen Tube-Synergid Interaction in Arabidopsis. Plant Physiology, 169(1), 485–496. 10.1104/pp.15.00528

Li, X., Bruckmann, A., Dresselhaus, T., & Begcy, K. (2024). Heat stress at the bicellular stage inhibits sperm cell development and transport into pollen tubes. Plant Physiology, 195(3), 2111–2128. 10.1093/plphys/kiae087

Mao, Y., Nakel, T., Erbasol Serbes, I., Joshi, S., Tekleyohans, D. G., Baum, T., & Groß-Hardt, R. (2023). ECS1 and ECS2 suppress polyspermy and the formation of haploid plants by promoting double fertilization. eLife, 12, e85832. 10.7554/eLife.85832

Mizuta, Y., Sakakibara, D., Nagahara, S., Kaneshiro, I., Nagae, T. T., Kurihara, D., & Higashiyama, T. (2024). Deep imaging reveals dynamics and signaling in one-to-one pollen tube guidance. EMBO reports, 25(6), 2529–2549. 10.1038/s44319-024-00151-4

Mòl, R., Matthys-Rochon, E., & Dumas, C. (1994). The kinetics of cytological events during double fertilization in Zea mays L. The Plant Journal, 5(2), 197–206. 10.1046/j.1365-313X.1994.05020197.x

Schindelin, J., Arganda-Carreras, I., Frise, E., Kaynig, V., Longair, M., Pietzsch, T., Preibisch, S., Rueden, C., Saalfeld, S., Schmid, B., Tinevez, J.-Y., White, D. J., Hartenstein, V., Eliceiri, K., Tomancak, P., & Cardona, A. (2012). Fiji : An open-source platform for biological-image analysis. Nature Methods, 9(7), 676–682. 10.1038/nmeth.2019

Sprunck, S. (2020). Twice the fun, double the trouble : Gamete interactions in flowering plants. Current Opinion in Plant Biology, 53, 106–116. 10.1016/j.pbi.2019.11.003

Sugi, N., Calhau, A. R. M., Jacquier, N. M. A., Millan-Blanquez, M., Becker, J. D., Begcy, K., Berger, F., Bousquet-Antonelli, C., Bouyer, D., Cai, G., Cheung, A. Y., Coimbra, S., Denninger, P., Dresselhaus, T., Feijó, J. A., Fowler, J. E., Geelen, D., Grossniklaus, U., Higashiyama, T., … Widiez, T. (2024). The peri-germ cell membrane : Poorly characterized but key interface for plant reproduction. Nature Plants, 10(11), 1607–1609. 10.1038/s41477-024-01818-5

Sugi, N., Izumi, R., Tomomi, S., Susaki, D., Kinoshita, T., & Maruyama, D. (2023). Removal of the endoplasma membrane upon sperm cell activation after pollen tube discharge. Frontiers in Plant Science, 14. 10.3389/fpls.2023.1116289

Wang, W., Malka, R., Lindemeier, M., Cyprys, P., Tiedemann, S., Sun, K., Zhang, X., Xiong, H., Sprunck, S., & Sun, M.-X. (2024). EGG CELL 1 contributes to egg-cell-dependent preferential fertilization in Arabidopsis. Nature Plants, 10(2), 268–282. 10.1038/s41477-023-01616-5

Zhong, S., Lan, Z., & Qu, L.-J. (2025). Ingenious Male–Female Communication Ensures Successful Double Fertilization in Angiosperms. Annual Review of Plant Biology, 76(Volume 76, 2025), 401–431. 10.1146/annurev-arplant-083123-071512

